# Aphasia severity mediates the relationship between attention and sentence comprehension

**DOI:** 10.1101/2025.09.26.678921

**Authors:** Emily J. Lenz, Arianna N. LaCroix

**Affiliations:** Department of Speech, Language, and Hearing Sciences, Purdue University, West Lafayette, IN

**Keywords:** aphasia, stroke, attention, orienting, executive control, sentence comprehension

## Abstract

**Background:** Attention deficits are common after stroke and exacerbate communication challenges in people with aphasia (PWA). Prior studies report mixed associations between attention and language, raising the possibility that these links are explained by aphasia severity. We examined whether alerting, orienting, and executive control attention differ by aphasia severity and whether severity mediates their relationship with sentence comprehension.

**Methods:** Fifty-eight PWA (latent, mild, moderate, severe) and 21 neurotypical controls completed the Attention Network Test, Western Aphasia Battery-Revised, and a sentence-picture matching task. Group differences were tested with ANOVA and linear mixed-effects models. Correlations and structural equation modeling evaluated associations among attention, aphasia severity, and sentence comprehension efficiency.

**Results:** Orienting and executive control differed by aphasia severity, with the severe group performing worse than all others; alerting did not differ across groups. Within group analyses showed that controls and mild aphasia participants demonstrated all three effects, latent aphasia participants showed orienting and executive control effects, while the moderate and severe groups only showed the executive control effect. Stronger orienting and executive control correlated with milder aphasia and more efficient canonical comprehension. Mediation confirmed that orienting influenced canonical and non-canonical comprehension indirectly through aphasia severity; executive control showed trend-level mediation. Alerting attention was unrelated to aphasia severity or comprehension.

**Conclusions:** Orienting and executive control, but not alerting, are closely tied to aphasia severity and indirectly support sentence comprehension. Attentional profiling may enhance individualized rehabilitation planning for PWA.

## Introduction

Aphasia affects approximately one-third of stroke survivors and creates major barriers to communication, independence, and quality of life. Beyond language impairments, up to 80% of people with aphasia (PWA) experience additional cognitive deficits in the first-year post-stroke (El Hachioui et al., 2014), among which attention difficulties are particularly consequential, as they contribute to poorer language performance (LaCroix, Blumenstein, et al., 2020; Murray et al., 1997; Murray, 2012; Varkanitsa et al., 2023; Villard & Kiran, 2017).

The attentional subsystems model defines attention as the coordinated function of three interacting networks: alerting (achieving and maintaining a state of readiness), orienting (shifting attention toward relevant stimuli), and executive control (resolving conflict among competing stimuli or responses while completing goal-directed behavior) (Petersen & Posner, 2012; Posner & Petersen, 1990). Although well-established in the neurotypical literature (de Souza Almeida et al., 2021), the subsystems model has only recently been applied to the study of attention in aphasia (Dash et al., 2020; Huang et al., 2022; LaCroix, Tully, et al., 2020; Mohapatra & Dash, 2023). Yet, this framework offers two potential advantages. First, the Attention Network Test efficiently assesses all three subsystems using a single cued-flanker task (Fan et al., 2002). Second, it is grounded in neuroanatomy, with each attentional subsystem being linked to relatively specific neural resources: alerting depends on a right-lateralized frontoparietal system and noradrenergic input from the locus coeruleus; orienting is supported by a bilateral dorsal frontoparietal network (including the intraparietal sulcus and frontal eye fields) together with a ventral network for reorienting; and executive control relies on the cingulo-opercular and dorsolateral prefrontal systems (Corbetta & Shulman, 2002; Markett et al., 2021; Petersen & Posner, 2012; Rinne et al., 2013; Xiao et al., 2016). Different lesion patterns across these networks may therefore yield distinct attentional deficit profiles in individual PWA.

Studies applying the subsystems model to aphasia have reported reduced efficiency in the alerting and orienting networks compared with controls (Dash et al., 2020; Huang et al., 2022; LaCroix, Tully, et al., 2020; Mohapatra & Dash, 2023). These findings align with earlier evidence that PWA have difficulty sustaining attention (Korda & Douglas, 1997; Laures, 2005) and orienting to salient stimuli (Petry et al., 1994; Robin & Rizzo, 1989; Villard & Kiran, 2015). Results for executive control are more mixed: some studies find no differences between PWA and controls (Dash et al., 2020; LaCroix, Tully, et al., 2020; Mohapatra & Dash, 2023), while others document impaired performance in PWA (Huang et al., 2022). The latter finding is consistent with the broader selective attention literature, which also reports executive control difficulties following a stroke (Erickson et al., 1996; Heuer & Hallowell, 2015; Murray, 2012; Pompon et al., 2015; Schumacher et al., 2022; Simic et al., 2020; Villard & Kiran, 2017).

Findings across studies highlight that the three attentional subsystems are differentially vulnerable in aphasia, but the functional consequences of these impairments for language remain less clear. Sentence comprehension tasks, particularly those contrasting canonical and non-canonical structures, provide one way to examine this link, as the more complex non-canonical sentences are thought to place greater demands on cognitive resources (e.g., Cutter et al., 2022; Just et al., 1996; Meng & Bader, 2025). Evidence from older neurotypical adults supports this view: reduced alertness in aging has been shown to compromise sentence processing (LaCroix, Blumenstein, et al., 2020), possibly because individuals are less prepared to process incoming speech. Such decreased readiness may also contribute to phonological, lexical, semantic, and repetition deficits in aphasia (Huang et al., 2022; Pérez Naranjo et al., 2018). Difficulty orienting to critical information such as the prosodic cues that mark clause structure and key words also disrupts sentence comprehension (LaCroix, Blumenstein, et al., 2020; Oakley, 2009; Schafer, 1997), and orienting deficits have additionally been linked to poorer repetition skills (Huang et al., 2022). Executive control and selective attention also play a crucial role, as difficulties suppressing distracting information result in poorer sentence comprehension, particularly in noisy environments (Fitzhugh et al., 2021; Huang et al., 2022; Kalbe et al., 2005; Villard & Kidd, 2019). Beyond comprehension, executive control deficits further contribute to conversational challenges (Frankel et al., 2007) and difficulties retelling stories in both quiet (Dutta et al., 2024) and noise (Harmon et al., 2024; Nelson et al., 2023).

Taken together, prior work indicates that variability in attentional processes may help explain the heterogeneity of language difficulties observed in PWA. At the same time, aphasia severity is a key determinant of this heterogeneity, as it captures the overall magnitude of impairment across language domains. In fact, severity is often the strongest predictor of post-stroke language outcomes, with greater severity linked to more extensive deficits and poorer treatment responses (Fridriksson & Hillis, 2021; Kristinsson et al., 2023; Nakagawa et al., 2019; Plowman et al., 2012). Despite this, relatively few studies have directly examined how attention relates to aphasia severity, and the results from existing studies are mixed: some report that poorer attention is associated with more severe aphasia and greater language impairment (Fonseca et al., 2019; Lee et al., 2020; Meier et al., 2022; Murray, 2012; Schumacher et al., 2022), whereas others find no relationship (Gordon-Pershey & Wadams, 2017; Murray et al., 1997; Yao et al., 2020).

Here, we examined whether the presence and extent of attention deficits varies as a function of aphasia severity. We also tested whether aphasia severity mediates the relationship between attention and language outcomes. We hypothesized that individuals with more severe aphasia would exhibit greater attention deficits than those with milder aphasia, both within- and between groups. Further, we predicted that aphasia severity would mediate the relationship between attention and language, such that weaker attention would be associated with poorer language outcomes as a consequence of greater aphasia severity.

## Method

### Participants

Fifty-eight left hemisphere stroke survivors participated in this study (24 female). Participants ranged in age from 31 to 82 years (*M* = 59.47, s*d* = 10.02), were pre-morbidly right-handed, >6 months post-stroke, and American English speakers who denied premorbid diagnosis of neurological disease, head trauma, or psychiatric diagnosis. Aphasia type and severity were determined using the *Western Aphasia Battery-Revised (*WAB-R; Kertesz, 2007). Overall, the sample included eight participants with latent aphasia (WAB-AQ >93.8), 23 with mild aphasia, 21 with moderate aphasia, 3 with severe aphasia, and 3 with very severe aphasia. We combined the severe and very severe aphasia participants into a single group due to their small sample sizes; we refer to this group as the “severe” aphasia group hereafter. Table 1 provides a detailed description of the stroke participants, including demographic and study-specific information.

**Table 1.** Participant demographic and study specific data.

| Subject | Gender (F/M) | Age | Race | Eth. | Ed. (Yr.) | TPS (Mo.) | WAB-AQ | Aphasia Type and Severity |
| --- | --- | --- | --- | --- | --- | --- | --- | --- |
| 1002 | F | 49 | White | Not H/L | 17 | 134 | 97.8 | Latent |
| 1013 | F | 60 | B/A | Not H/L | 21 | 82 | 96.8 | Latent |
| 1015 | F | 43 | White | Not H/L | 17 | 27 | 95.6 | Latent |
| 24 | F | 55 | White | H/L | 16 | 98 | 95.5 | Latent |
| 1007 | M | 58 | White | Not H/L | 17 | 31 | 95.5 | Latent |
| 1039 | M | 76 | White | Not H/L | 17 | 96 | 95.4 | Latent |
| 1018 | M | 66 | White | Not H/L | 21 | 45 | 95.1 | Latent |
| 6 | M | 44 | White | NR | 16 | 63 | 95.0 | Latent |
| 17 | M | 65 | White | Not H/L | 20 | 107 | 93.2 | Mild Anomic |
| 14 | F | 57 | White | Not H/L | 18 | 67 | 93.0 | Mild Anomic |
| 1 | M | 31 | NR | H/L | 14 | 52 | 93.0 | Mild Anomic |
| 1019 | F | 56 | B/A | Not H/L | 19 | 438 | 92.8 | Mild Anomic |
| 1041 | M | 67 | White | Not H/L | 19 | 37 | 92.0 | Mild Anomic |
| 1009 | M | 63 | White | Not H/L | 19 | 146 | 91.9 | Mild Anomic |
| 1014 | F | 51 | White | Not H/L | 19 | 281 | 89.3 | Mild Anomic |
| 27 | M | 59 | White | Not H/L | 12.5 | 6 | 88.6 | Mild Anomic |
| 30 | F | 72 | White | H/L | 9 | 81 | 88.1 | Mild Anomic |
| 1045 | F | 49 | White | Not H/L | 17.5 | 116 | 87.1 | Mild Anomic |
| 1033 | F | 50 | White | Not H/L | 17 | 47 | 87.0 | Mild Anomic |
| 1010 | M | 58 | White | H/L | 21 | 46 | 85.7 | Mild Anomic |
| 1040 | F | 63 | White | Not H/L | 19 | 62 | 85.6 | Mild Anomic |
| 1038 | M | 55 | B/A | Not H/L | 17 | 68 | 84.8 | Mild Anomic |
| 1035 | M | 59 | White | Not H/L | 17 | 86 | 84.1 | Mild Anomic |
| 2 | F | 49 | White | Not H/L | 12 | 91 | 83.8 | Mild Anomic |
| 1025 | M | 61 | Asian | Not H/L | 21 | 137 | 82.7 | Mild Anomic |
| 1006 | M | 63 | White | Not H/L | 17 | 72 | 81.5 | Mild Conduction |
| 1030 | M | 71 | White | Not H/L | 19 | 80 | 80.6 | Mild Anomic |
| 1029 | F | 61 | White | Not H/L | 19 | 33 | 79.2 | Mild Broca's |
| 1028 | M | 62 | White | Not H/L | 19 | 237 | 78.9 | Mild Anomic |
| 1047 | M | 72 | White | Not H/L | 13 | 179 | 77.2 | Mild Anomic |
| 1037 | M | 67 | B/A | Not H/L | 19 | 36 | 76.2 | Mild Anomic |
| 1042 | M | 59 | B/A | Not H/L | 13 | 61 | 74.9 | Moderate Broca's |
| 1043 | M | 54 | White | Not H/L | 17 | 94 | 74.6 | Moderate Anomic |
| 1023 | F | 61 | Asian | Not H/L | 19 | 104 | 74.2 | Moderate Wernicke's |
| 1022 | M | 70 | White | Not H/L | 17 | 35 | 74.2 | Moderate Transcortical Motor |
| 1001 | F | 52 | White | Not H/L | 17 | 84 | 73.8 | Moderate Conduction |
| 31 | F | 77 | White | Not H/L | 12 | 120 | 73.2 | Moderate Conduction |
| 1052 | F | 45 | White | Not H/L | 17 | 97 | 70.7 | Moderate Conduction |
| 1044 | M | 57 | White | Not H/L | 19 | 87 | 70.3 | Moderate Conduction |
| 1012 | M | 82 | White | Not H/L | 19 | 39 | 70.0 | Moderate Conduction |
| 25 | F | 49 | White | Not H/L | 12 | 20 | 69.9 | Moderate Transcortical Motor |
| 1032 | M | 46 | White | Not H/L | 17 | 96 | 67.3 | Moderate Broca's |
| 9 | M | 63 | White | H/L | 14 | 176 | 64.9 | Moderate Broca's |
| 12 | F | 73 | White | H/L | 18 | 74 | 64.7 | Moderate Broca's |
| 1011 | M | 65 | White | Not H/L | 15 | 60 | 63.2 | Moderate Broca's |
| 1036 | M | 67 | White | Not H/L | 17 | 49 | 62.8 | Moderate Broca's |
| 1021 | M | 46 | White | Not H/L | 17 | 42 | 62.0 | Moderate Broca's |
| 1046 | M | 70 | White | Not H/L | 13 | 43 | 61.8 | Moderate Wernicke's |
| 32 | F | 65 | White | Not H/L | 12 | 27 | 59.7 | Moderate Transcortical Motor |
| 1053 | F | 41 | White | Not H/L | 19 | 39 | 59.4 | Moderate Broca's |
| 18 | F | 49 | White | Not H/L | 16 | 31 | 55.9 | Moderate Broca's |
| 1016 | M | 74 | White | Not H/L | 13 | 50 | 52.2 | Moderate Wernicke's |
| 1027 | M | 66 | White | Not H/L | 17 | 35 | 37.7 | Severe Broca's |
| 1034 | M | 59 | White | Not H/L | 15 | 103 | 36.8 | Severe Broca's |
| 29 | M | 59 | White | Not H/L | 16 | 13 | 36.1 | Severe Broca's |
| 22 | F | 58 | White | Not H/L | 18 | 129 | 23.6 | Very Severe Broca's |
| 1024 | M | 66 | White | Not H/L | 16 | 73 | 17.9 | Very Severe Broca's |
| 28 | F | 64 | White | Not H/L | 15.5 | 7 | 16.3 | Very Severe Broca's |
| PWA<br>Avg. | 24/34 | 59.47 | --- | --- | 16.71 | 83.85 | 74.50 | --- |
| Control<br>Avg. | 10/21 | 57.14 | --- | --- | 15.62 | --- | --- | --- |

Note. F = female; M = male; Eth. = ethnicity; Ed. = education; Yr. = years; TPS = time post-stroke; Mo = months; WAB-AQ = Western Aphasia Battery–Revised Aphasia Quotient; H/L = Hispanic/Latino; B/A = Black or African American; NR = no response; Avg. = average scores across all participants in each group.

Twenty-one right-handed, American English speakers served as the control group (10 female). Control participants ranged in age from 24 to 77 years (*M* = 57.14, s*d* = 13.04) and were matched to the stroke group on age and education (*p* >.05). All participants provided written informed consent prior to participating. Study procedures were approved by the Midwestern University (IRB AZ1464) and Purdue University (IRB 2023-1067) Institutional Review Boards.

### Procedure

This study is a retrospective analysis combining data from two previous studies involving left hemisphere stroke participants who completed the WAB-R, Attention Network Test (ANT), and a sentence-picture matching task. The original studies, conducted at Purdue University and Midwestern University, examined the effects of music on attention in PWA (Purdue) and the relationship between attention and language in PWA (Midwestern). Forty-one PWA completed the Purdue study, while 17 PWA and the control group completed the procedures as part of the Midwestern University study.

#### Attention Network Test (ANT)

Figure 1 includes a schematic of an ANT trial, as well as examples of the cue conditions and targets (Fan et al., 2002, 2009). Each trial began with a fixation cross jittered between 500-2000 (Midwestern) and 2400-3600 (Purdue) milliseconds. Following the offset of the fixation cross, participants were presented with a cue condition for 100 milliseconds (Figure 1B). Cue conditions for both studies included double cue (i.e., two asterisks presented simultaneously above and below the fixation cross), spatial cue (i.e., single asterisk presented above or below the fixation cross; spatial cues predicted the location of the upcoming flanker task with 75% accuracy), and no cue (i.e., no asterisk presented; fixation cross remained in the center of the screen). Participants in the Purdue study also saw a center cue (i.e., single asterisk presented in the center of the screen), however this condition was excluded from the analyses since it was not present in the Midwestern paradigm. Following the offset of the cue, participants were presented with a second fixation cross for 400 milliseconds, then the flanker task.

**Figure 1.**
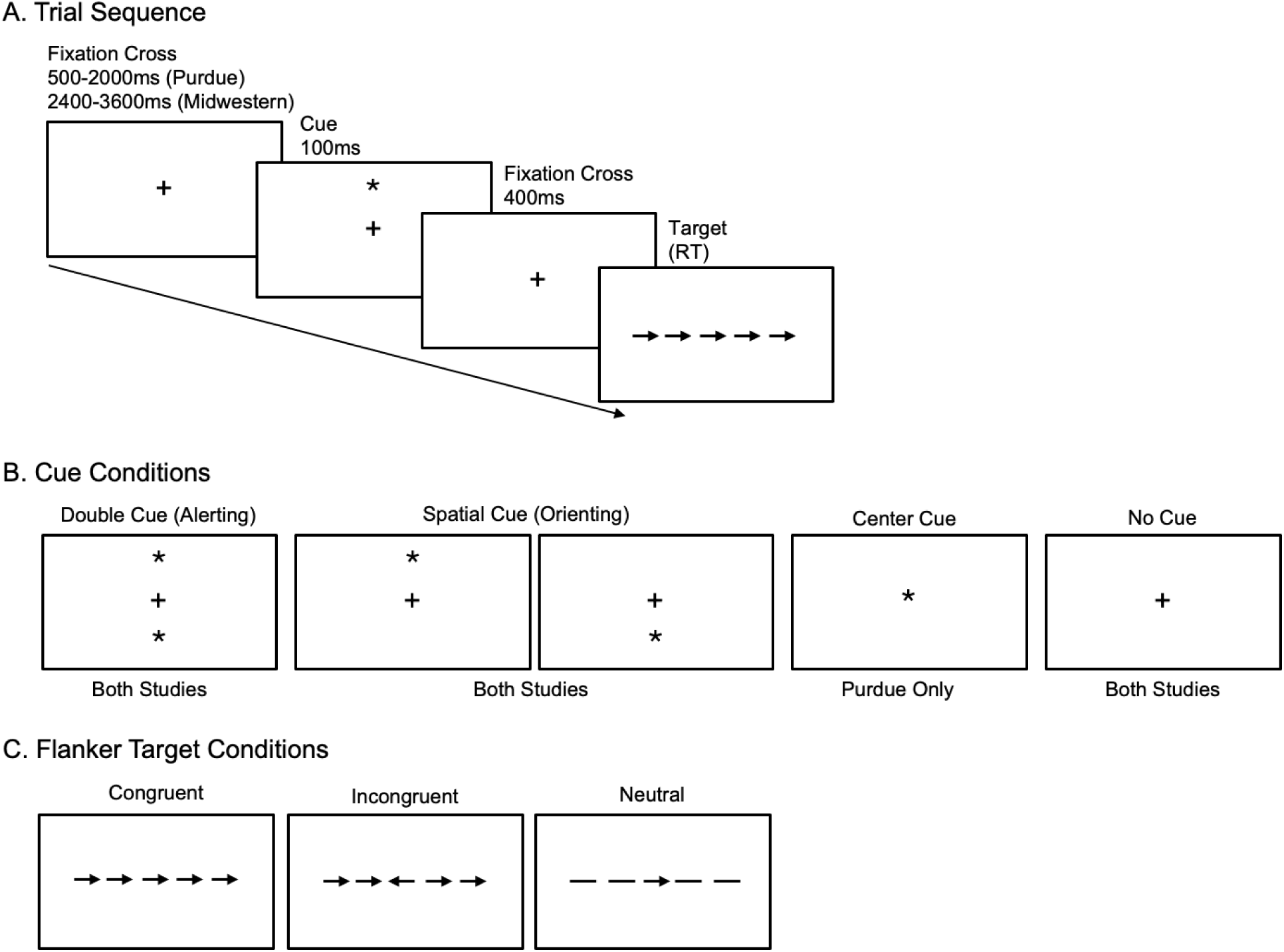
Attention Network Test trial sequence (A), cue conditions (B), and flanker targets (C).

In each flanker trial, participants were presented with a series of five stimuli consisting of a central target arrow flanked by either congruent arrows that face the same direction, incongruent arrows that face the opposite direction, or neutral flankers (two horizontal lines on each side) (Figure 1C). Participants were instructed to select which direction (left or right) the central arrow faced as quickly and accurately as possible. Reaction time and accuracy were recorded for each trial via keyboard button presses. To account for potential right-sided hemiparesis in the aphasia group, all participants responded using their left hand. The “z” and “x” keys were labeled with left (<) and right (>) arrow stickers, respectively, to reinforce left-hand responding.

Participants received verbal and written instructions and completed 5 practice trials prior to the onset of the experimental task. Purdue participants completed 156 experimental trials, while Midwestern participants completed 70 experimental trials. In line with the ANT literature (e.g., Fan et al., 2002), the efficiency of the alerting subsystem was computed by comparing reaction times on no cue and double cue trials, orienting by comparing reaction times on double cue and spatial cue trials, and executive control by comparing reaction times on incongruent and neutral trials.

#### Sentence-Picture Matching Task

The sentence stimuli have been described in detail in previous work (LaCroix, Blumenstein, et al., 2020; LaCroix et al., 2019; Wilson et al., 2010). Briefly, participants listened to two sentence structures spoken with a natural speech prosody. Each sentence was comprised of 10 syllables, two nouns (boy, girl), one of seven verbs (kick, wash, chase, push, kiss, pull, hug), and one of three colors (blue, green, red). The canonical sentences had a subject-relative structure (e.g., The boy who is blue is chasing the girl) and the non-canonical sentences had an object-relative structure (e.g., The girl who the boy is chasing is blue). The key difference between these two syntactic structures is that subject-relatives preserve English’s canonical subject-verb-object order within the embedded relative clause, whereas object-relatives disrupt this order within the relative clause by fronting the object (represented by the relative pronoun *who* for *the girl*), resulting in an apparent object-subject-verb sequence.

Each trial began with the simultaneous presentation of a binaurally presented sentence and two pictures (one target, one foil). Participants were instructed to decide which picture matched the sentence as quickly and accurately as possible. Reaction time and accuracy were collected for each trial via keyboard press (Purdue: “z” and “x” keys; Midwestern: “f” and “j” keys). Verbal and written instructions preceded the start of the task. Participants in the Purdue study completed 5 practice trials followed by 70 experimental trials, of which 34 (17 canonical and 17 non-canonical) were included in the analysis. Participants in the Midwestern study completed 3 practice trials followed by 80 experimental trials, of which 20 (10 canonical and 10 non-canonical) were included in analyses.

### Statistical Analyses

All data were analyzed using RStudio version 4.4.1 (R Core Team, 2025). All ANT trials (156 from Purdue and 70 from Midwestern) were included in the analyses, as restricting the Purdue dataset to the first 70 ANT trials per participant did not qualitatively change the results. Reaction times (RTs) for incorrect responses and those exceeding 2.5 standard deviations from a participant’s mean were excluded to ensure that the analyses reflected the attentional processes of interest rather than artifacts of distraction or motor response errors (e.g., Thériault et al., 2024). Following this approach, 2.41% of trials were excluded for controls (.004% errors, 2.41% long RTs) and 1.62% of trials for PWA (.02% errors, 1.60% long RTs). The remaining trials, consisting of correct responses only, were used in the analyses, as accuracy was high in both groups (98.18 ± 13.36% across PWA and 98.69 ± 11.39% in controls).

#### Impact of Aphasia Severity on Attention: Between Group Differences

From the remaining reaction time data, we computed RT difference scores for each participant as follows: alerting (no cue – double cue), orienting (double cue – spatial cue), and executive control (incongruent – neutral trials). These difference scores are consistent with the ANT literature (Fan et al., 2002), and are designed to isolate attention-specific effects from individual differences in overall reaction time or processing speed, thereby enabling more meaningful between-group comparisons. We conducted three separate ANOVAs, one for each attentional subsystem. In all models, the dependent variable was the RT difference score, and the independent variable was aphasia severity (control, latent, mild, moderate, severe).

#### Impact of Aphasia Severity on Attention: Within Group Effects

We additionally investigated whether the alerting (no cue vs. double cue), orienting (double cue vs. spatial cue), and executive control (incongruent vs. neutral trials) effects were present *within* each group. To do this, we computed a linear mixed effects model using the lmer function from the lme4 package (Bates et al., 2015). The dependent variable was the log-transformed reaction time, which was applied to meet model assumptions of variance homogeneity. The fixed effects were cue (no cue, double cue, spatial cue), congruency (congruent, incongruent, neutral), aphasia severity (control, latent, mild, moderate, severe), and study site (Purdue, Midwestern). Aphasia severity was categorical and based on the WAB-R Aphasia Quotient (WAB-AQ). We also specified the cue x aphasia severity and congruency x aphasia severity interactions, as these would allow us to investigate the presence of each effect *within* each group. Subject was specified as a random intercept to account for individual variability in response times. The results were first summarized as an Analysis of Deviance using car::Anova (Fox & Weisberg, 2019). Pairwise comparisons for the fixed and interaction effects were computed using “emmeans” (Length, 2023). Multiple corrections were corrected using the Benjamini-Hochberg (BH) procedure (Benjamini & Hochberg, 1995).

#### Relationship Between Attention, Aphasia Severity, and Language

Only participants with aphasia were included in the following analyses. Pearson correlations were specified with cor.test in base R (R Core Team, 2025). Alerting, orienting, and executive control attention were each represented by the RT difference score and separately correlated with aphasia severity (WAB-AQ), each subtest of the WAB-R (spontaneous speech, auditory comprehension, repetition, naming/word finding), and canonical and non-canonical sentence comprehension. Sentence comprehension was operationalized using a Balanced Integration Score (BIS), which combines accuracy and response time into a single efficiency metric robust against speed accuracy tradeoffs (Liesefeld & Janczyk, 2019). Higher BIS scores reflect more efficient sentence processing.

Finally, we tested whether aphasia severity mediated the relationship between attention and sentence comprehension using structural equation modeling implemented in lavaan (Rosseel, 2012). The three attention measures (RT difference scores for alerting, orienting, and executive control) were specified as exogenous predictors, and aphasia severity (indexed by the WAB-AQ) was included as the mediator. Sentence comprehension, operationalized as BIS efficiency scores for canonical and non-canonical sentences, served as the outcome variable. The WAB-R subtests were excluded from this model because they contribute to the WAB-AQ. All direct paths were specified, and the residual variances of canonical and non-canonical BIS scores were allowed to covary to account for shared method variance. Standard errors were estimated using nonparametric bootstrapping with 5,000 draws. Both unstandardized and standardized path coefficients, along with 95% bootstrap confidence intervals, were reported.

## Results

### Impact of Aphasia Severity on Attention: Between Group Differences

The overall ANOVA models demonstrated that orienting, F(4, 74) = 4.70, *p =* .002, and executive control attention, F(4, 74) = 14.19, *p <* .001, differed by aphasia severity, but that alerting attention did not, F(4, 74) = .95, *p =* .44. All pairwise comparisons are reported in Table 2 and graphed in Figure 2. The severe aphasia group had poorer orienting (Figure 2B) and executive control attention (Figure 2C) than all other groups, but no other group differences were observed.

**Table 2.** Pairwise comparisons of alerting, orienting, and executive control attention across different levels of aphasia severity.

| Comparison | Alerting | Orienting | Executive Control |
| --- | --- | --- | --- |
| Control vs. Latent | $t = 1.05, p = .59$ | $t = -1.37, p = .25$ | $t = -.20, p = .85$ |
| Control vs. Mild | $t = -.30, p = .77$ | $t = -.63, p = .53$ | $t = -.89, p = .63$ |
| Control vs. Moderate | $t = .80, p = .61$ | $t = .91, p = .41$ | $t = -1.05, p = .60$ |
| Control vs. Severe | $t = .80, p = .61$ | $t = 3.33, p = .005^*$ | $t = -7.23, p < .001^*$ |
| Latent vs. Mild | $t = -1.29, p = .59$ | $t = .92, p = .41$ | $t = -.45, p = .82$ |
| Latent vs. Moderate | $t = -.46, p = .72$ | $t = 2.04, p = .09$ | $t = -.58, p = .80$ |
| Latent vs. Severe | $t = -1.57, p = .59$ | $t = 3.91, p = .002^*$ | $t = -6.05, p < .001^*$ |
| Mild vs. Moderate | $t = 1.12, p = .59$ | $t = 1.56, p = .21$ | $t = -.18, p = .85$ |
| Mild vs. Severe | $t = -.70, p = .61$ | $t = 3.77, p = .002^*$ | $t = -6.72, p < .001^*$ |
| Moderate vs. Severe | $t = -1.42, p = .59$ | $t = 2.72, p = .02^*$ | $t = -6.53, p < .001^*$ |

\*Significant $p < .05$ . Degrees of freedom equal 74 for all pairwise comparisons.

**Figure 2.**
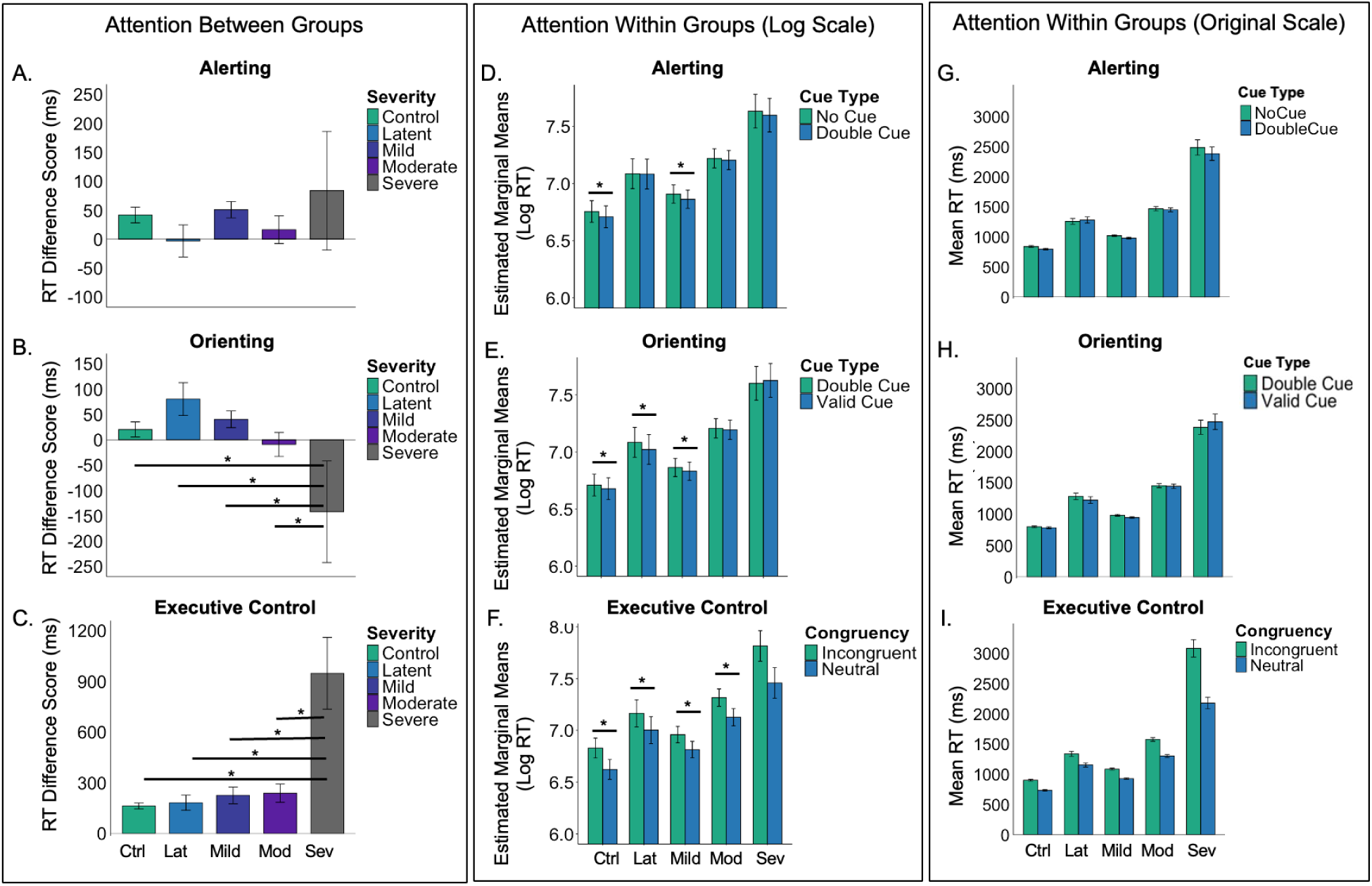
The left panel shows between-group differences in attention, represented as RT difference scores for each subsystem: (A) alerting = no cue – double cue, (B) orienting = double cue – spatial cue, and (C) executive control = incongruent – congruent targets. Larger positive values indicate faster responses to the alerting (double) cue, orienting (spatial) cue, and neutral flanker target. The middle panel presents estimated marginal means showing comparisons for attention within each group: (D) alerting (no cue vs. double cue), (E) orienting (double cue vs. spatial cue), and (F) executive control (incongruent vs. neutral). The right panel presents mean RTs showing comparisons for attention within each group: (G) alerting, (H) orienting, and (I) executive control. Error bars represent ±1 SD. Note. ms = milliseconds, Ctrl = control, Lat = latent, Mod = moderate, Sev = severe.

### Impact of Aphasia Severity on Attention: Within Group Effects

The full model results are reported in Table 3. The fixed effect of cue and congruency were both significant. As expected, all participants were slower on the no cue than double cue trials (the alerting effect), double cue than spatial cue trials (the orienting effect), and incongruent compared to neutral trials (the executive control effect). The fixed effect of aphasia severity^1^ was also significant, as were the aphasia severity x cue and aphasia severity x congruency interactions, which allowed us to explore the presence of each attentional effect within each group. As reported in Table 4 and depicted in Figure 2D-I, the control group demonstrated all three effects in the expected direction (i.e., better performance on double cue, spatial cue, and congruent trials), as did the mild aphasia group. However, the latent aphasia group only demonstrated the orienting and executive control effects, but not the alerting effect, whereas the moderate and severe aphasia groups demonstrated the executive control effect only.

**Table 3.** Full model results for the LMER model.

| Effect | Statistic |
| --- | --- |
| Intercept | $F(1, 73.1) = 17050.57, p < .001^*$ |
| Congruency | $F(2, 6829.3) = 340.61, p < .001^*$ |
| Cue Condition | $F(2, 6829.1) = 2.43, p < .001^*$ |
| Aphasia Severity | $F(4, 73.0) = 9.5, p < .001^*$ |
| Study Site | $F(1, 73.1) = 19.32, p = .123$ |
| Congruency x Aphasia Severity | $F(8, 6829.2) = 6.91, p < .001^*$ |
| Cue Condition x Aphasia Severity | $F(8, 6829.1) = 2.22, p = .023^*$ |
| *Significant $p < .05$ | |

**Table 4.** Pairwise comparisons showing the significance of the alerting, orienting, and executive control effects within each group.

|  | Alerting | Orienting | Executive Control |
| --- | --- | --- | --- |
| Control | $t = 3.03, p = .005^*$ | $t = 2.07, p = .04^*$ | $t = 13.35, p < .001^*$ |
| Latent | $t = .17, p = .87$ | $t = 2.93, p = .007^*$ | $t = 7.68, p < .001^*$ |
| Mild | $t = 3.62, p = .001^*$ | $t = 2.57, p = .01^*$ | $t = 11.58, p < .001^*$ |
| Moderate | $t = 1.11, p = .33$ | $t = .97, p = .33$ | $t = 14.23, p < .001^*$ |
| Severe | $t = 1.29, p = .33$ | $t = -.98, p = .33$ | $t = 13.17, p < .001^*$ |
| *Significant $p < .05$ | | | |

### Relationship Between Attention, Aphasia Severity, and Language

#### Correlations

Alerting attention was negatively correlated with the efficiency of canonical sentence comprehension, indicating that PWA who responded more quickly to the double cue showed less efficient canonical sentence processing (Table 5). Alerting attention did not correlate with aphasia severity (Figure 3, left panel), or any other language measure. Orienting attention was positively correlated with aphasia severity (Figure 3, center panel), all four WAB-R subtests, and canonical sentence comprehension efficiency, such that faster responses to the spatial cue were associated with less severe aphasia, milder language deficits overall, and more efficient canonical sentence processing. Executive control was negatively correlated with aphasia severity (Figure 3, right panel), all four WAB-R subtests, and canonical sentence comprehension efficiency, indicating that better executive control was linked to less severe aphasia, milder language deficits, and more efficient canonical sentence processing. Non-canonical sentence comprehension efficiency did not correlate with any attention measure.

**Table 5.**
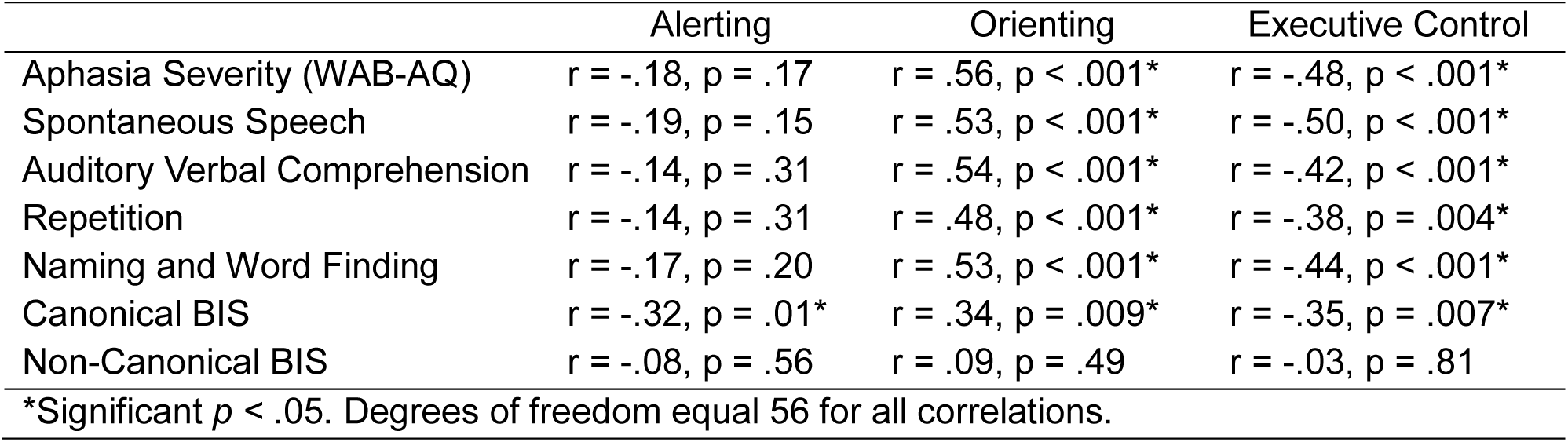
Pearson correlations between each attentional subsystem and language variable.

**Figure 3.**
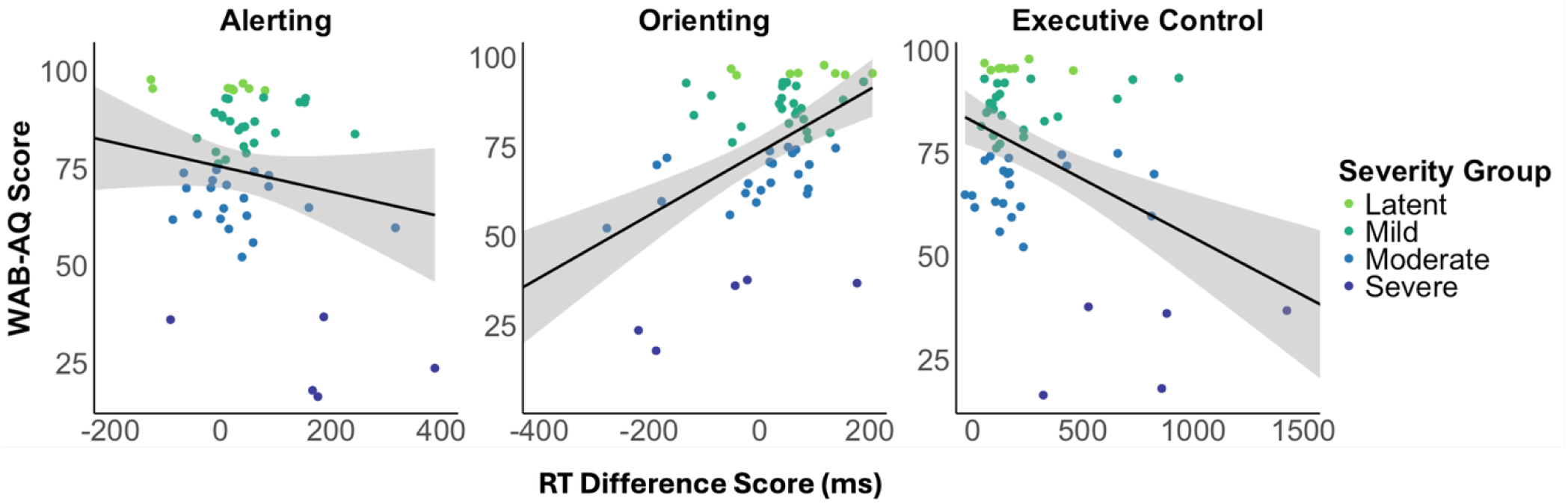
Scatterplots depicting the correlations between each attentional subsystem and aphasia severity (WAB-AQ). Larger more positive scores equate to faster performance following the alerting (double) cue, orienting (spatial) cue, and neutral flanker targets. Confidence intervals are +/- 95%.

#### Structural Equation Model

The full model results are presented in Supplementary Table 2 and illustrated in Figure 4. Orienting (β = .49, *p* = .001) and executive control (β = –.38, *p* = .019) predicted aphasia severity, such that stronger attention was associated with milder aphasia. In turn, aphasia severity predicted both canonical (β = .58, *p* = .003) and non-canonical efficiency (β = .51, *p* = .012), with milder aphasia linked to more efficient sentence processing. The direct effects from alerting, orienting, and executive control to sentence comprehension efficiency were all non-significant. The indirect effects confirmed that orienting attention influenced both canonical (β = .28, *p* = .019) and non-canonical sentence comprehension efficiency (β = .25, *p* = .032) through aphasia severity, while executive control showed trend-level indirect effects (p < .10). Collectively, the mediation analysis indicates that the effect of orienting attention on sentence processing in PWA is fully mediated by aphasia severity. In other words, orienting supports comprehension indirectly by reducing aphasia severity, rather than exerting a direct influence on sentence processing efficiency.

**Figure 4.**
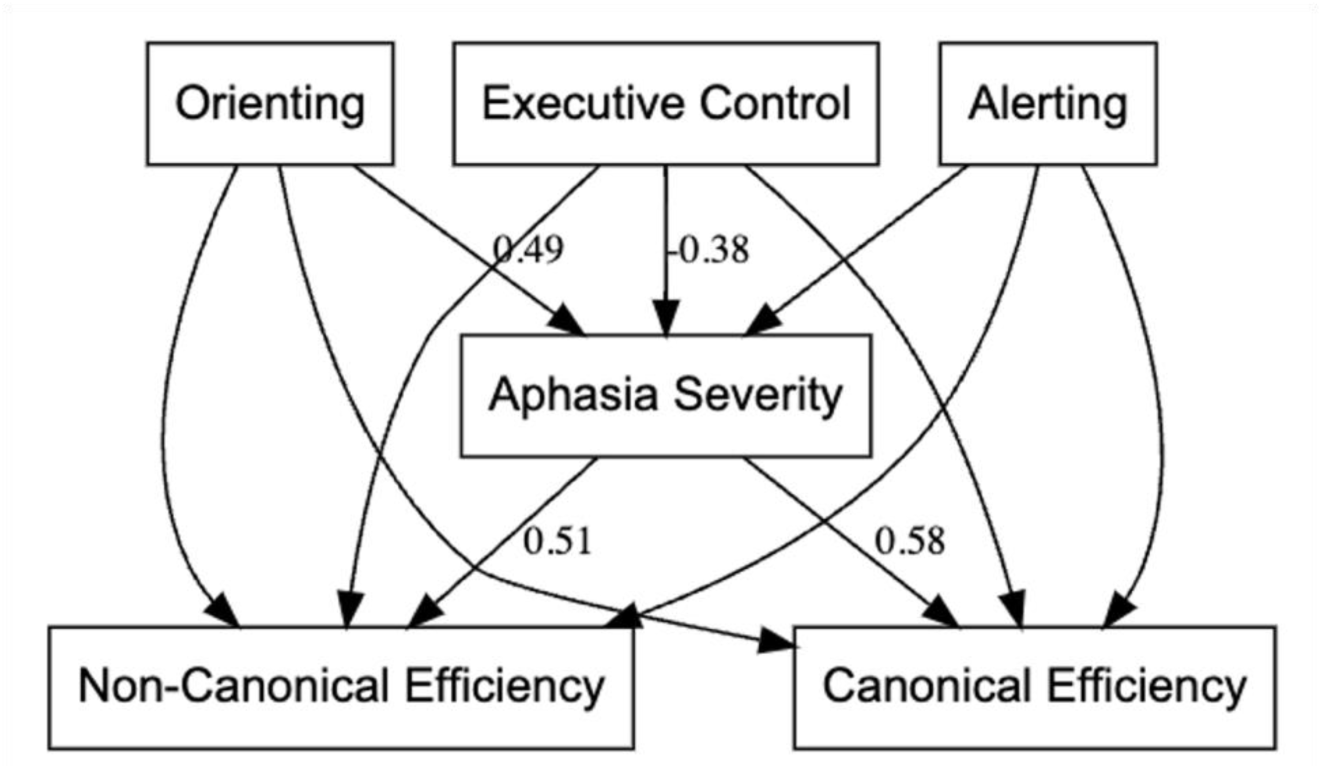
Mediation model illustrating the relationships among the attentional subsystems, aphasia severity, and sentence comprehension efficiency. All model paths tested are displayed; only standardized coefficients significant at *p* < .10 are labeled. Better orienting and executive control were associated with less severe aphasia, which in turn predicted greater efficiency for both canonical and non-canonical sentence comprehension. Alerting was not significantly related to aphasia severity or sentence comprehension efficiency.

## Discussion

The relationship between attention and language in aphasia is not always clear, with some studies reporting associations (Dutta et al., 2024; Fitzhugh et al., 2021; Frankel et al., 2007; Harmon et al., 2024; Huang et al., 2022; Kalbe et al., 2005; LaCroix, Blumenstein, et al., 2020; Nelson et al., 2023; Pérez Naranjo et al., 2018; Villard & Kidd, 2019) and others finding none (Gordon-Pershey & Wadams, 2017; Murray et al., 1997; Yao et al., 2020). Aphasia severity, however, is consistently linked to both attention and language (Fonseca et al., 2019; Lee et al., 2020; Meier et al., 2022; Murray, 2012; Schumacher et al., 2022), suggesting it may be a key factor linking the two processes, which may explain the mixed findings. To this end, we examined how attention deficits vary as a function of aphasia severity and tested whether severity mediates the relationship between attention and language.

### Attention and Aphasia Severity

Alerting attention did not differ significantly across groups and showed no relationship with aphasia severity. The absence of an association with severity is consistent with previous findings (Huang et al., 2022), yet the lack of group differences contrasts with prior reports of reduced alerting in PWA compared to controls (Dash et al., 2020; Korda & Douglas, 1997; LaCroix et al., 2020; Laures, 2005; Mohapatra & Dash, 2023). While these group-level results suggest that alerting may be relatively preserved in aphasia and not systematically tied to severity, the within-group analyses provide additional nuance: the alerting effect (double < no cue) was only significant in the control and mild aphasia groups.

The absence of an alerting effect in the moderate and severe groups may reflect diaschisis, whereby left hemisphere lesions disrupt the right-lateralized alerting network or subcortical structures (e.g., the thalamus) that support alerting across both hemispheres. The absent alerting effect in the latent aphasia group may also be explained by subcortical damage, however, this damage is likely direct to structures such as the thalamus (Fritsch et al., 2022). Alternatively, generalized slowing could explain the absent alerting effect in these three groups, particularly since response times were markedly slower in all three groups compared to the control group. Such slowing may reduce sensitivity to phasic alerting cues, which operate on a relatively brief timescale (<500 ms) (Fan et al., 2002; Hackley, 2009). In other words, when baseline processing speed is substantially delayed, participants may be unable to take advantage of the transient benefit provided by an alerting cue, even if the underlying neural mechanisms for alerting remain partially intact. This account highlights the importance of considering general processing speed when evaluating attention in aphasia, as slowed responses may mask or diminish the appearance of otherwise preserved attentional effects.

An emerging body of research has documented orienting attention impairments in post-stroke aphasia (Huang et al., 2022; LaCroix, Tully, et al., 2020; Petry et al., 1994; Robin & Rizzo, 1989; Villard & Kiran, 2015). The present findings are consistent with this literature and extend it by showing that orienting deficits scale with aphasia severity. Specifically, the orienting effect (spatial < double) was absent in the moderate and severe groups but present in the latent and mild aphasia groups. Moreover, most participants with severe aphasia (5/6) responded more slowly to the spatial cue than to the double cue, suggesting that the spatial cue may actually interfere with performance—a reversal previously observed in accuracy (LaCroix, Tully, et al., 2020). By contrast, most participants with latent aphasia (6/8) demonstrated the expected orienting effect, highlighting that orienting is preserved when lesions are smaller or more localized (and/or possibly subcortical) but increasingly compromised with greater lesion extent, particularly when the frontal and parietal cortices are involved.

The executive control effect (incongruent < neutral) was significant in all groups and correlated positively with aphasia severity, such that individuals with more severe aphasia showed disproportionately larger interference costs. This pattern supports prior evidence that executive control deficits worsen with increasing severity (Huang et al., 2022; Meier et al., 2022; Murray, 2012). Importantly, only the severe group differed from controls, which may help reconcile mixed findings in the literature. Studies reporting no group differences often sampled primarily from mild-to-moderate participants (Dash et al., 2020; LaCroix et al., 2020; Mohapatra & Dash, 2023), whereas those identifying deficits included more individuals with severe aphasia (Huang et al., 2022). These findings suggest that executive control deficits are not necessarily a uniform feature of aphasia, but emerge as aphasia severity increases, underscoring the importance of considering severity when characterizing domain-general cognitive impairments in this population.

### Attention, Language, and Aphasia Severity

The mediation model showed that the link between attention and sentence comprehension (both canonical and non-canonical) was explained by aphasia severity. Specifically, orienting and executive control predicted severity: individuals who oriented more efficiently to spatial cues and who were less distracted by incongruent flankers tended to have less severe aphasia. In turn, reduced severity was associated with more efficient sentence comprehension. For orienting, this pathway extended to both canonical and non-canonical sentences, whereas for executive control the relationship with comprehension was only a trend (*p <* .10). These findings extend prior evidence that orienting and executive attention support language in post-stroke aphasia by showing that aphasia severity mediates this relationship.

Smaller alerting effects (i.e., similar performance across no cue and double cue trials) were associated with more efficient canonical sentence comprehension. Although faster responses are typically interpreted as reflecting a greater benefit from alerting cues (Fan et al., 2002), a smaller alerting effect may instead signal more stable baseline attentional readiness. Individuals who maintain high tonic alertness may not require external cues to become prepared to process the sentence as they are, in effect, already ready.

Alerting attention was not significant in any path of the mediation model, despite its correlation with canonical sentence comprehension. This suggests that while alerting and comprehension co-vary, their relationship is not explained by aphasia severity, nor does alerting exert a unique direct effect once severity is included. One possibility is that the correlation reflects individual differences in attentional strategy: participants with more efficient comprehension may rely on sustained tonic alertness rather than transient cue-driven phasic alerting. In this view, alerting attention functions as a background or modulatory influence, supporting comprehension indirectly through stable readiness rather than directly through rapid cue processing. The processing speed results lend some credibility to this account but confirming it will require future studies that directly measure tonic alertness through sustained attention tasks or physiological indices such as pupil diameter.

### Clinical Implications

The mediation results suggest that attention does not directly drive sentence comprehension but instead influences it indirectly through aphasia severity. Of the three attention networks, orienting and executive control emerged as the most consequential. Alerting attention, by contrast, was less systematically related to the language impairment, potentially allowing it to function more as a background support mechanism.

The severity gradient may provide further clinical guidance. For instance, in latent and mild aphasia, alerting and orienting are relatively preserved and executive control deficits are subtle. Patients at these severity levels may still benefit from preparatory or spatial cues to support comprehension, and interventions that strengthen executive control could further bolster language processing. In moderate aphasia, alerting and orienting effects are diminished and executive control costs are elevated. These patients may struggle to capitalize on external cues and to manage competing linguistic demands. Therapy may therefore need to simplify environments, reduce distraction, and provide structured scaffolding. In severe aphasia, all three networks are compromised: alerting cues provide little benefit, orienting cues may even interfere with performance, and executive control deficits are most pronounced. Here, therapy may need to minimize reliance on flexible attention, instead emphasizing slower pacing, predictable routines, and reduced cognitive load.

### Limitations and Future Directions

While informative, a few limitations should be acknowledged. First, the sample size within individual severity groups, particularly latent and severe aphasia, was relatively small. This limits statistical power and may have contributed to variability in effects such as the absence of an alerting effect in latent aphasia. Future research with larger samples is needed to confirm the severity-dependent patterns observed here.

Second, to enhance statistical power and ensure representation across aphasia severities, we combined data from two related studies. This approach increased the sample size but introduced a minor methodological difference: participants completed slightly different numbers of experimental trials on the ANT and sentence-picture matching task across studies. Because the tasks and stimuli were otherwise identical, and no differences emerged in accuracy or reaction time, this variation is unlikely to have influenced the findings. Nevertheless, it should be considered when interpreting the results, and future research should seek to replicate these effects using a single standardized protocol.

Lesion data was not collected as part of this study. Although medical records provided some insight into neural damage, the lack of neuroimaging prevented us from directly linking attention deficits to lesion location or network disruption. Incorporating structural and functional imaging in future work will be critical for understanding the mechanisms that may contribute to aphasia-related attentional impairments.

Finally, the mediation analysis showed that orienting and executive control influence comprehension primarily through aphasia severity. However, the mechanisms underlying this pathway and the exacerbating factors remain unclear. Future work with larger samples and more fine-grained measures will be needed to determine whether particular linguistic abilities (e.g., lexical retrieval, syntactic processing), cognitive–linguistic interactions, or individual factors influence this mediation.

### Conclusions

This study shows that attentional profiles in aphasia seem to vary with severity, and that the influence of orienting and executive control attention on sentence comprehension is largely mediated by the overall impairment. These severity-dependent patterns help reconcile mixed findings in the literature and underscore severity as a central link between attention and language. Clinically, the results suggest that strengthening orienting and executive control may indirectly improve comprehension, while therapy for severe aphasia may need to emphasize more compensatory strategies.

## Footnotes

1 The fixed effect of aphasia severity was significant. However, this effect likely reflects processing speed rather than attention, as it captures overall reaction time differences across groups rather than group differences based on the difference scores isolating each attentional subsystem, as reported in Table 2. Pairwise comparisons exploring this fixed effect are reported in Supplementary Table 1. Overall, the control group was faster than the latent aphasia group, the mild aphasia group was faster than the moderate aphasia group, and the moderate aphasia group was faster than the severe aphasia group. The latent aphasia group did not differ from the mild aphasia group.

